# An insight into transcriptome of *LlaNAC* Gene Over-expressing Tobacco Plants

**DOI:** 10.1101/2020.06.24.169250

**Authors:** Sadhana Singh, Atul Grover

## Abstract

Here, we report a whole transcriptome analysis of *LlaNAC* gene (from *Lepidium latifolium*) containing transgenic tobacco line (NC10) and wild type (WT), to attain deeper knowledge into the downstream genes activated by the over-expressing transgene. Transcriptome sequencing of NC10 and WT samples generated huge data using Illumina platform. The maximum number of unigenes GO annotated were of Biological process (8988, 3209) followed by molecular function (5155, 2577) and cellular components (3826, 1583) for WT and NC10 samples respectively. KEGG Pathway analysis revealed the unigenes were enriched in different functional pathway categories. The unigenes whose products involved in carbohydrate metabolism, glycan metabolism, and secondary metabolites synthesis were more for NC10 library in comparison to WT. Greater variety of transcription factors were involved in transgenic than wild-type plants. Genes like, Copia-like retrotransposable element, Peroxidase 64-like, Peptidyl-prolyl cis-trans isomerise, Cytochrome P450, Lipoyl synthase, CBL-interacting serine/threonine-protein kinase 5-like etc. were found differentially expressed in both the samples. Promoter analysis of these differentially expressed genes have elements for defence and stress response, abscisic acid response, shoot specific expression and light response, etc. In summary this study reports the involvement of the overexpressed genes in the dual action of cold tolerance and biomass accumulation, as sugars participate in both of these activities of the cell.

## 1. Introduction

About 10% of the genes encodes transcription factors (TFs) of plant transcriptome, which typically function as immediate or early stress responsive genes [1,2]. TFs regulate gene expression by binding to cis-acting elements in the promoters of the target genes encoding proteins with specific functions under varied biological context [3]. Some of these TFs are central regulators of signalling and regulatory pathways of stress adaptation. A number of transcription factors, such as CBF/DREB, NAC, zinc finger protein, bZIP, MYB, WRKY, APETALA, basic helix-loop-helix (bHLH), C2-H2 type zinc fingers (ZFs), etc. are known to play significant roles in translating abiotic stress signals to changes in gene expression thereby affecting overall metabolomic environment of the cell [4,1]. A typical transcription factor contains a DNA-binding region, an oligomerization site, a transcription-regulation domain and a nuclear localization signal (NLS).

NAC is one of the largest TF families in plants and in most plants the copy number of NAC genes are known to be more than 70 [2]. The acronym NAC finds its origin to three different genes (*NAM, ATAF, CUC*) into which the NAC domain was first reported. NAC family TFs contain a highly conserved DNA binding NAC domain (150 amino acids) in N-terminal and a diversified C-terminal domain that generally regulates transcriptional activation [5,6,7,8]. NAC TFs have a variety of important functions in plant development, morphogenesis, senescence, and abiotic stress responses [9,10,11,2,12,13]. In recent years, as more and more NAC genes from different plants are being functionally characterized, the knowledge on diversity of the functions performed is being enriched [14,15,16]. We have earlier reported a NAC gene from *Lepidium latifolium* (*LlaNAC*), whose over-expression was found directly correlated with the accumulation of biomass in *Nicotiana tabacum* (tobacco) transgenic lines. In addition, these lines were shown to mature early, had shorter life cycles and were more tolerant to abiotic stresses and sequestered more carbon than non-transformed plants [9,8]. We had also carried out in silico assessments to suggest possible mechanisms for these phenotypic effects [9]. Here, we report a whole transcriptome analysis of one of these *LlaNAC* containing tobacco lines vis-a-vis wild type (WT), to gain an insight into the downstream genes activated by the over-expressing transgene (*LlaNAC*).

## 2. Results and discussion

### 2.1 Validation of Transgenic Lines

More than 90% seeds on an average of all the transgenic lines successfully germinated under limiting concentration of paromomycin (150 ppm), implying that all the lines had achieved homozygosity. A minor fraction of seeds (∼5%) did not germinate, which could be due to their individual viability. Line NC10 showed 100 percent survival rate. PCR based validation of grown up plants was carried out using the primer pair LlaNAC-TqF (5’-ACA GTG GTA AAC CTC CAA AAG G-3’) and LlaNAC-TqR (5’-CGA AGA GAG TTC TTG TTG ACG A-3’) to obtain a band of size 122 bp. Upon dual validation (by germination under limiting concentration of antibiotics and PCR assay), plants were assessed for various morphometric traits (Table S1). Though role of NAC transcription factors in secondary growth of plants has been known for quite some time (reviewed by Singh et al. [13], their direct involvement in biomass production has been a relatively recent observation [9,17]. Besides that, the T3 generation of tobacco over-expressor plants continues to show overall changes in life cycle, consistent with our findings on T2 generation [9]. Real time-based quantitative assays were carried out for these transgenic lines at 50 and 100 DAS. Large amount of transcript accumulation was observed in case of NC10 line, i.e. nearly 1,800-fold increase on 100 DAS as compared to WT plants. Whereas approximately 700-fold elevation could be seen in rest of the three lines i.e. NC2, NC18 and NC7b (Table S1). Earlier, Ordiz et al. [18] also reported more than 450-fold induction of gene expression in transgenic tobacco plants containing the zinc finger (TFs ZF) protein and ß-glucuronidase reporter gene construct.

### 2.2 Impact of LlaNAC Gene on Transcriptome of Transformed Plants

When a gene is introduced into a new genome, it is likely to make many changes to the transcriptomic environment and thereby in the metabolic environment of the cell. Such changes may also occur due to the site of the integration of the gene, as the site itself may have epistatic and other positional effects [19]. In cases, where a transgene is a transcription factor, the effects are manifold as the encoded protein would bind to a number of diverse genes which would induce the expression of the other genes. Transformation of tobacco with the *LlaNAC* gene has thus been expected to produce varied phenotypic effects and altered physiological responses [8]. We earlier reported that *LlaNAC* led to enhancement of biomass, shortened life cycle, early maturity and cold stress tolerance in tobacco [9]. Thus, it has become all the more important to carefully assess what particular genes are being affected by the transformation.

We attempted to identify the transcriptomic make-up of the transgenic plants, by sequencing of the whole transcriptome. Total RNA was isolated from the transgenic (NC10) and WT plants and run on 1% denatured agarose gel. High quality data was generated on the Illumina platform (NextSeq). Trimmomatic v 0.30 was used for filtering of raw data. At the first place, having comparable yields and quality of total RNA, three times more data was generated for the transgenic sample compared to WT. The high-quality reads were assembled using Trinity. A total of 95,799 Unigenes spanning 50,008,638 bases and 51,379 Unigenes spanning 26,187,994 were assembled for NC10 and WT samples, respectively. While, it may be a direct consequence of large amount of reads created for the NC10 sample, it may also be possible due to expression of more number of genes in the transgenic compared to the WT. Only a small subset of Unigenes (7,176) were common to the two libraries. Interestingly, 74% of the Unigenes from NC10 had blast hits, while only 66% of Unigenes from WT sample had a blast hit. Conversely, almost 25,000 Unigenes from the NC10 library had not been reported earlier, and 17,500 Unigenes from WT library had not been reported earlier. Thus, together they constitute a huge collection of sequences for which function is still required to be associated, thereby creating a huge reserve for gene discovery. The average length of Unigenes in NC10 sample were 522 bases, with longest Unigene being 18,561 bases long, while the average length of Unigenes from WT sample was 510 bases and longest being 12,439 bases long. Nevertheless, gene annotations of sequences with known homologs, mainly mapped down to metabolic pathways of major biomolecules (Table 1).

**Table 1.** Pathway predictions of Unigenes based on mining of KEGG database.

| Description | NC10 sample | WT sample |
| --- | --- | --- |
| **Metabolism** |  |  |
| Carbon metabolism | 242 | 134 |
| Carbohydrate metabolism | 395 | 167 |
| Energy metabolism | 230 | 151 |
| Lipid metabolism | 206 | 86 |
| Nucleotide metabolism | 108 | 69 |
| Amino acid metabolism | 259 | 127 |
| Metabolism of other amino acids | 130 | 48 |
| Glycan biosynthesis and metabolism | 98 | 31 |
| Metabolism of cofactors and vitamins | 133 | 52 |
| Metabolism of terpenoids and polyketides | 122 | 49 |
| Biosynthesis of other secondary metabolites | 115 | 40 |
| Xenobiotics biodegradation and metabolism | 42 | 19 |
| **Genetic information processing** |  |  |
| Transcription | 116 | 78 |
| Translation | 353 | 260 |
| Folding, sorting and degradation | 272 | 146 |
| Replication and repair | 123 | 75 |
| **Environmental information processing** |  |  |
| Signal transduction | 321 | 146 |
| Signaling molecules and interaction | 2 | 2 |
| **Cellular processes** |  |  |
| Transport and catabolism | 203 | 78 |
| Cell motility | 30 | 14 |
| Cell growth and death | 131 | 85 |
| Cellular community | 46 | 17 |
| **Organismal systems** |  |  |
| Environmental adaptation | 78 | 31 |

The functional annotation for the Unigenes was carried out by aligning them to non-redundant database of NCBI using blastx (cut off E-value 1e-06). The majority of the hits were found against tobacco, as expected. The Unigenes GO annotated for both the samples were for Biological process (3209 for NC10 sample, and 8988 for WT sample) followed by molecular function (2577 for NC10 sample, and 5155 for WT sample) and cellular components (1583 for NC10 sample, and 3826 for WT sample), as indicated in Figure 3. Ortholog assignment and mapping of the Unigenes to the biological pathways were performed using KEGG automatic annotation server (KAAS). All the Unigenes were compared against the KEGG database using blastx with threshold bit-score value of 60 (default). The mapped Unigenes represented metabolic pathways of major biomolecules such as carbohydrates, lipids, nucleotides, amino acids, glycans, cofactors, vitamins, terpenoids, polyketides, etc. The mapped Unigenes also represented the genes involved in genetic information processing, environmental information processing and cellular processes (Table 1). On fine assessment of these Unigenes, it was found that nearly 19% of the Unigenes whose products were involved in metabolism had roles in carbohydrate metabolism, 4.7% were involved in glycan metabolism and 5.52% in synthesis of secondary metabolites in case of NC10 library. In comparison, in WT library these ratios stood at 17.16, 3.19 and 4.11 percents respectively. Further, nearly twice the number of genes involved in signal transduction in response to environmental information could be mapped in case of the transgenic as compared to WT (Table 1). The Unigenes were also compared against the COG database. Surprisingly, more matches were obtained in WT sample compared to the NC10 sample. For NC10 sample, 4,896 Unigenes had significant homology, while for WT sample 5,284 Unigenes had significant homology (Figure 1). These Unigenes could be assigned to 24 functional categories.

**Figure 1.**
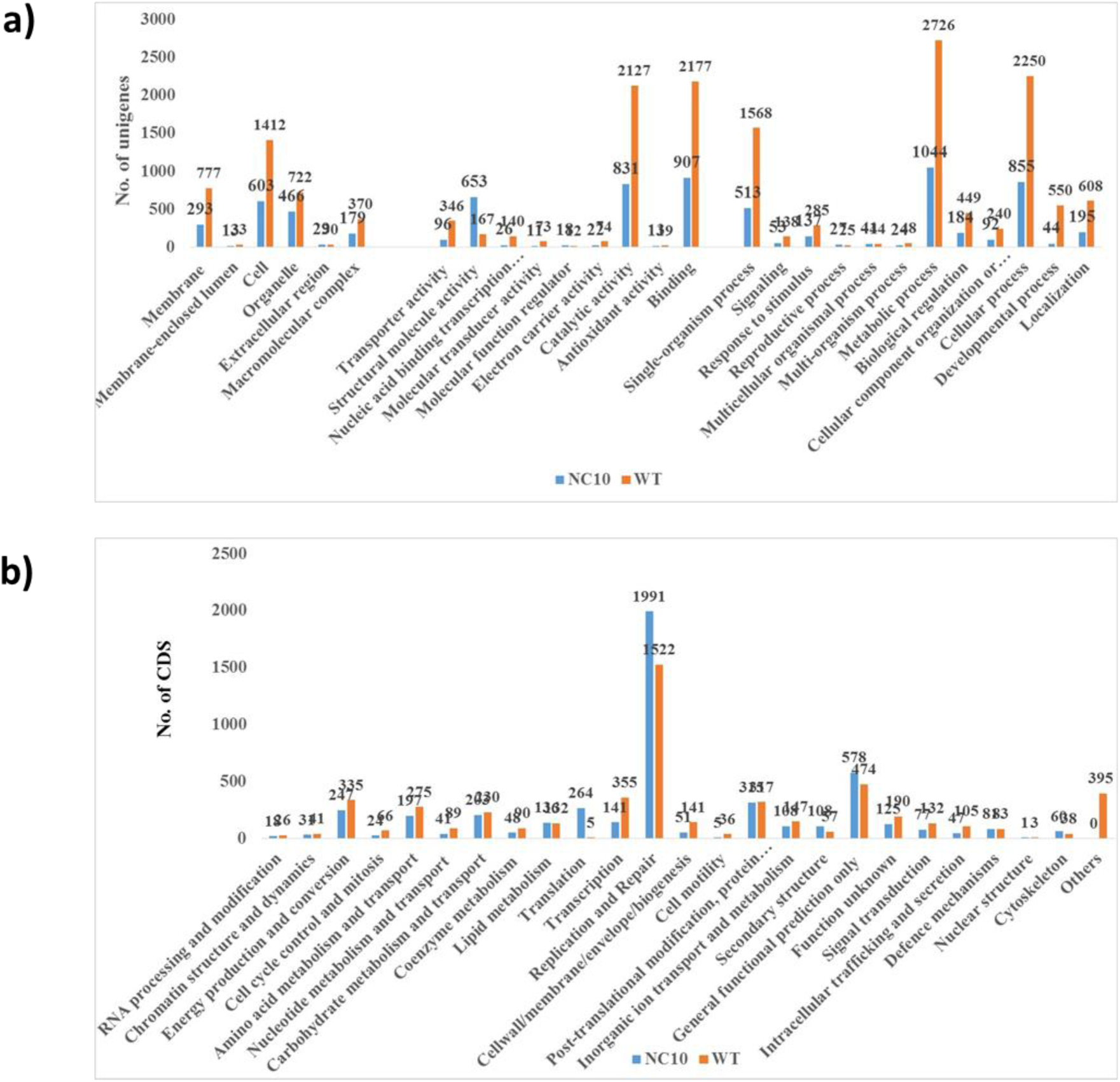
**(a**) Comparative GO functional classifications, i.e., Cellular Component, Molecular Function and Biological Processes for both the NC10 and WT samples **(b)** COG classification of unigenes from NC10 and WT sample

Transcription factors were identified by search against the Plant transcription factor database (PlnTFDB; http://plntfdb.bio.uni-potsdam.de/v3.0/downloads.php) using BLASTX with an E-value cut-off of < 1e-05. The maximum Unigenes had aligned against FAR1 TFs family (Figure 2). Therefore, it may be safely concluded that more number of target genes are being induced for expression in the transgenic plants compared to the WT. For the genes related to transcription including transcription factors, though more genes were found expressed in WT, there were greater variety of genes involved in case of the transgenic, and hence possibly affecting the expression of more genes in the genome, which in turn might be inducing multiple characters of agronomic importance to the plants.

**Figure 2.**
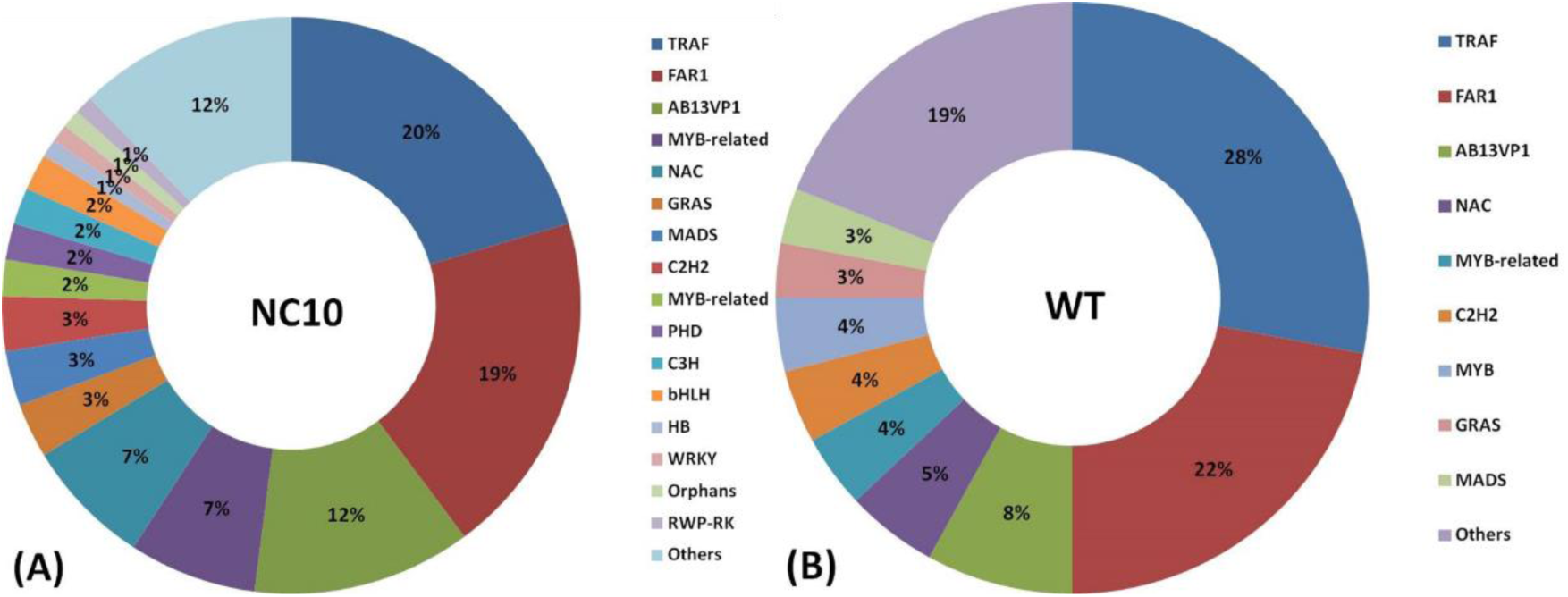
Transcription family distribution of (A) NC10 and (B) WT samples

Further, we found several CAREs such as, cis-acting element for high transcription levels (5UTR Py-rich stretch), abscisic acid responsive (ABRE), anerobic induction (ARE), heat responsive element (HSE), common element in promoter and enhancer region (CAAT-box), gibberellin responsive element (GARE-motif, P-box), auxin responsive (AuxRR core), salt responsive element (GT1-motif), core promoter element around −30 of transcription start site (TATA-box), zein metabolism regulation (O2-site), endosperm expression (Skn-1_motif, GCN4), special protein 1 motif (Sp1), defence and stress response (TC-rich repeats), salicylic acid response (TCA-element), light responsive (TCT-motif, I-box, GA-motif, G-box, AE-box, GAG-motif), circadian control (circadian), shoot specific expression and light responsive (as-2-box) etc. indicating that these elements have role in building biomass and might play major role in conferring tolerance against various type of environmental stresses (Table S2).

Genes like Organelle transcript processing protein, ATP-dependent DNA helicase PIF1, Copia-like retrotransposable element, Endonuclease/reverse transcriptase, ABC transporter G family member 29-like, Transcription factor TGA2, Cytochrome P450 83B1-like, Peroxidase 64-like, Lipoyl synthase (chloroplastic), CBL-interacting serine/threonine-protein kinase 5-like etc. were found differentially expressed in NC10 sample and WT sample based on the p-value significance (**Table 2**). Further, prediction of subcellular localization based on Plant-mPLoc database [20] (http://www.csbio.sjtu.edu.cn/bioinf/plant-multi/) revealed that most of the differentially expressed genes were predominantly localized in the chloroplast, and nucleus while some of them were located in the peroxisome, cell membrane, mitochondria, vacuole, and endoplasmic reticulum. However, few DEGs were having multiple localization sites.

**Table 2.**
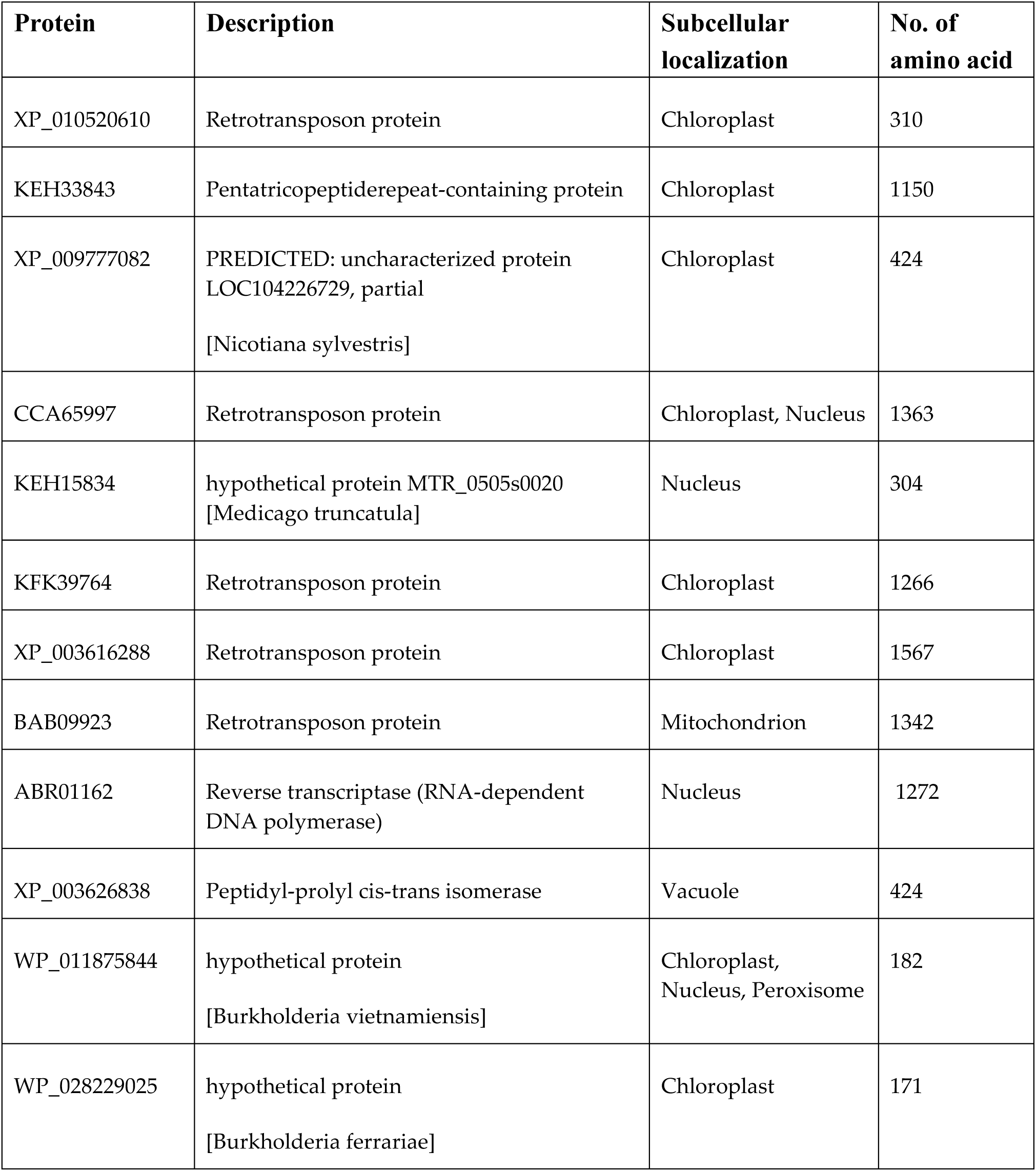
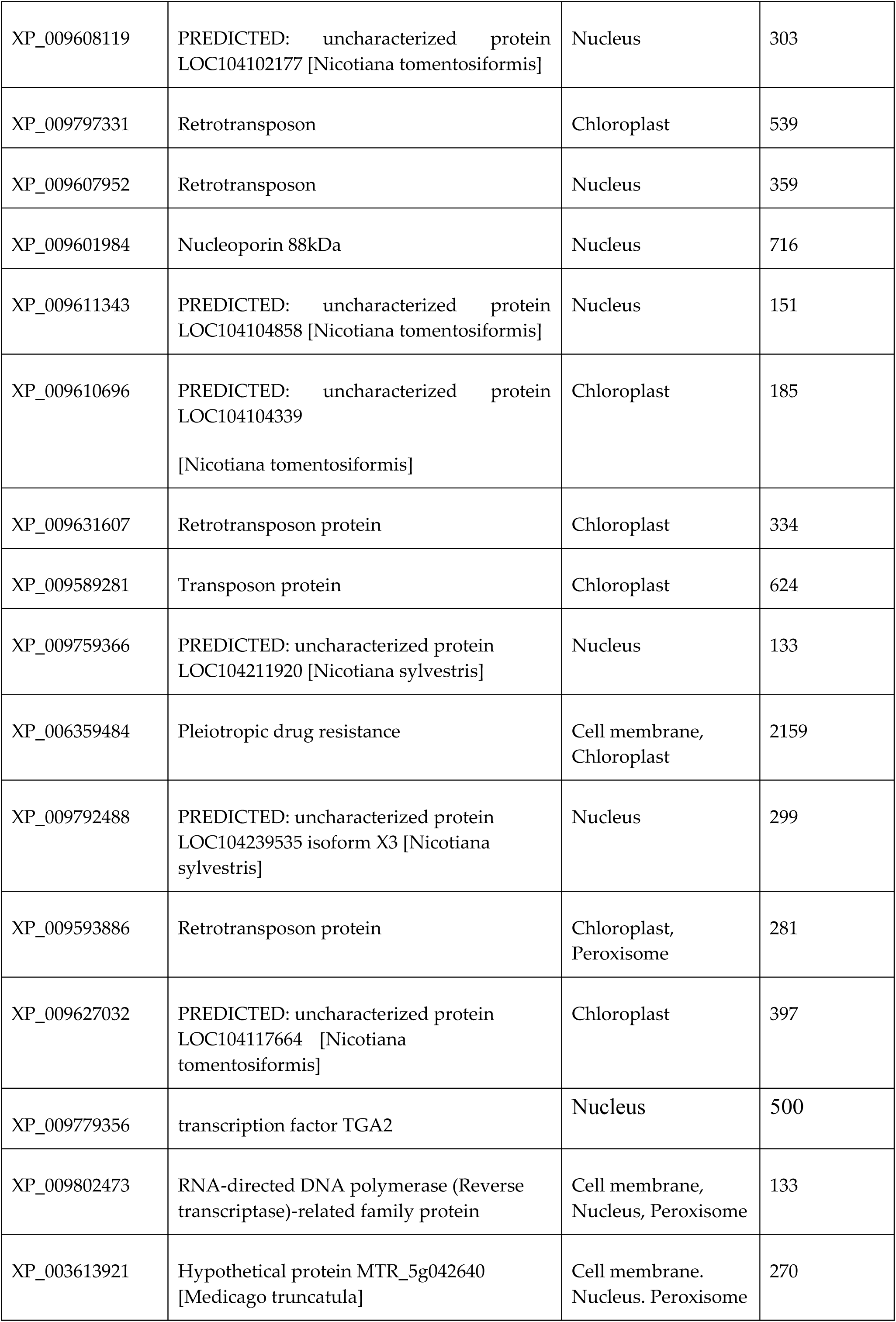
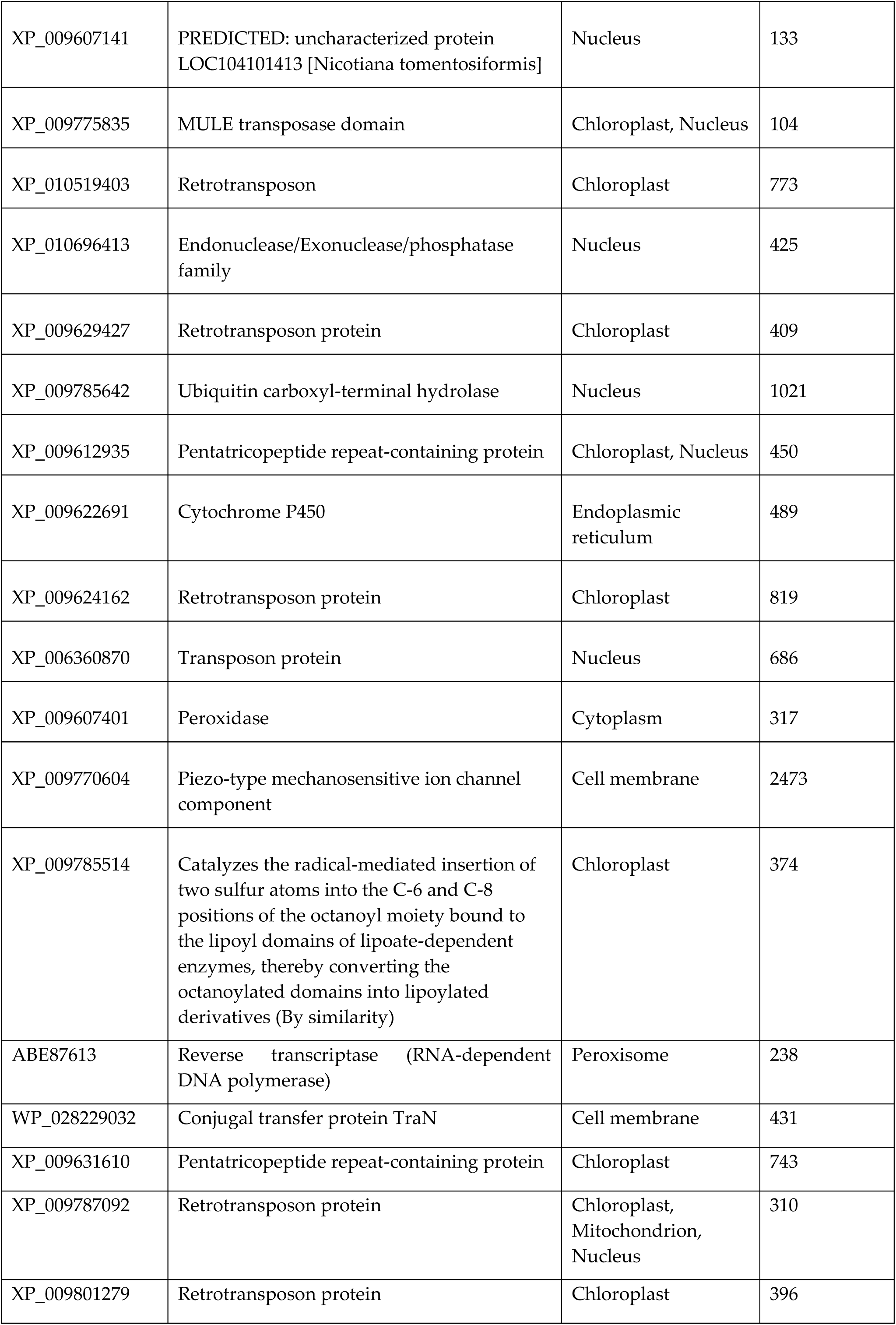
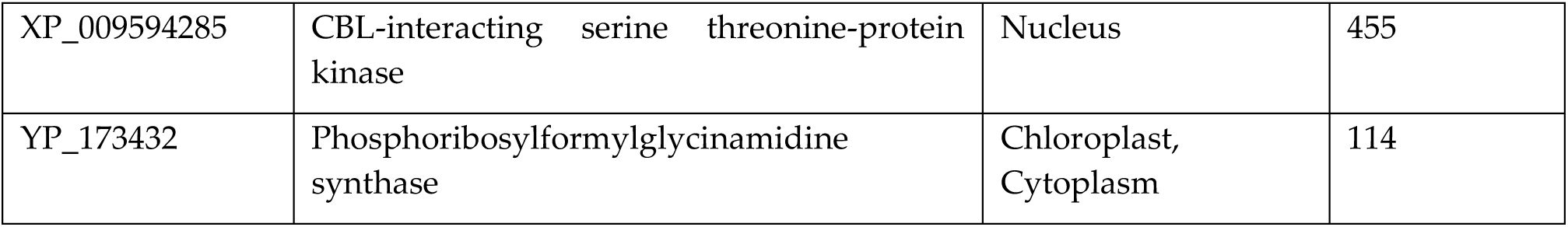
List of common Unigenes between NC10 and WT found differentially expressed based on the p-value significance and their subcellular localization prediction based on Plant-mPLoc database

## 4. Materials and Methods

### 4.1 Plant material, Growth conditions and Validation of Genetically Modified Lines

T3 seeds *Nicotiana tabacum* (tobacco) plants over-expressing *LlaNAC* gene, and co-transformed *npt*<u>II</u> gene, (four transgenic lines-NC2, NC7b, NC10 and NC18 in generations T3) along with wild-type (WT) plants were grown in antibiotic containing moist filter paper (150 ppm) under controlled conditions of temperature (25 ± 2°C) and light (16/8 h photoperiod). Seedlings germinated upto the cotyledonary leaf stage was recognized as the ‘survival event’ in the presence of the antibiotic. After 15 days of germination, seedlings were transferred to soilrite for further experimental analysis. The plants were nourished with MS basal medium twice a week, and were watered (distilled water) on daily basis. Each individual plant was tested for its genetic stability based on its tolerance to 150 ppm paromomycin, following PCR assay with an initial denaturation at 94°C for 5 min., followed by 30 cycles of 94°C for 30 s and 55°C for 30 s and 72°C for 1.5 min. and a final extension of 10 min. at 72°C. Amplicons thus obtained were run on 1% agarose gel to check the presence or absence of *LlaNAC* gene. We have conducted the analysis on T3 generation, and therefore, characterization of the growth-related parameters is also presented in the supplementary Table (S1). Growth parameters e.g., leaf area, plant height, midrib length was recorded at the 50 days and 100 days of sowing (DAS) of seeds. Total RNA was isolated from leaf samples using HiPurA™ Plant RNA isolation Kit (HiMedia laboratories Pvt. Ltd, India) following instructions in user manual. Equal amount (1.0 g) of the RNA quantified using Qubit fluorometer (Invitrogen, USA) was used for first-strand cDNA synthesis using QuantiTect Reverse Transcription kit (QIAGEN, Germany) following manufacturer’s instructions. Real time PCR was carried out with *LlaNAC* primers using cDNA as template for NAC transcript abundance at 50 and 100 DAS.

### 4.2 Global Analysis of Transcriptome in NAC Transgenic Line NC10

The analysis of transcriptome of a NAC transgenic plant was carried out by harvesting the leaf tissue from three plants of about 30 days and pooling them. NC10 was chosen as the representative line. A similar procedure was followed with the WT plants. Total RNA was isolated from using Xcelgen Plant RNA Isolation Kit (Xceleris Genomics, India) as per the manufacturer’s protocol. Library preparation, sequencing and analysis was outsourced to M/s Xceleris, India. Reverse transcription of the total RNA to complementary DNA and amplification of cDNA templates by long-distance PCR (LD-PCR) were performed using SMARTer Ultra-low Input RNA for Illumina Sequencing-HV (Clontech Labaratories, CA, USA) kit. The paired-end cDNA sequencing libraries were prepared using Illumina TruSeq Nano DNA HT Library Preparation Kit as per the described protocol.

### 4.3 Functional Annotation

The functional annotation was performed by aligning Unigenes those to nr database of NCBI using BLASTX with an E-value threshold of 1e-06. GO assignments were used to classify the functions of the Unigenes. The GO mapping also provided ontology of defined terms grouped into three main domains: Biological process, Molecular function and Cellular component. Ortholog assignment and mapping of the Unigenes to the biological pathways were performed using KEGG automatic annotation server (KAAS). All the Unigenes were compared against the KEGG database using BLASTX with threshold bit-score value of 60 (default). The Unigenes were compared against the COG database for the analysis for widespread domain families. For the identification of transcription factor families, the Unigenes of NC10 and WT samples were searched against all the transcription factor protein sequences at Plant transcription factor database [21] (PlnTFDB; http://plntfdb.bio.uni-potsdam.de/v3.0/downloads.php) using BLASTX with an E-value cut-off of < 1e-05.

### 4.4 Promoter Identification

To investigate transcriptional regulation of these differentially expressed genes, 1500 bp upstream sequences of translational start site were analysed using PlantCARE database [22] for the presence of cis-acting regulatory elements (CAREs).

## 5. Conclusions

*LlaNAC* is an interesting gene, which was identified from a cold induced subtraction library [23]. It has subsequently been introduced in tobacco in this laboratory, and the over-expressor line has shown a variety of interesting characters related to the growth, life cycle and stress tolerance [9]. Objective of the present study had been to assess the possible mechanism by which such diversity of effects was being produced. Whole transcriptome sequencing is a powerful tool to assess the overall cellular environment against a control. Being a transcription factor, *LlaNAC* was expected to affect a number of downstream genes, and the same has been recorded by us earlier as well [9]. Many of these downstream genes themselves are transcription factor, and these in turn affect further genes within the genome. This partially explains the diversity of effects due to a singly transformed gene. Further, as majority of the over-expressed genes in the dataset were mapped to sugar metabolism pathway, it became a strong indication for its dual action of cold stress tolerance and biomass accumulation, as sugars participate in both these activities of the cell [24,25,26,27,28,29]. Changes in the life cycle and growth patterns are complicated traits. NAC genes are known to be associated with development and growth [30,31,32], but the present data has not been sufficient to explain the transcriptomic changes leading to shortening of the life cycle of the plants. This research is parallel to the ageing trait in animals, and more dedicated research excursions are required to understand its total biology.

## Supplementary Materials

Supplementary materials can be found at www.mdpi.com/xxx/s1.

## Author Contributions

**S**.S. conceived the idea and performed the experiment/analysis. S.S. and A.G. together wrote the MS.

## Acknowledgments

Sadhana Singh acknowledged the fellowship received during research work from DRDO.

---

**Table S1.**
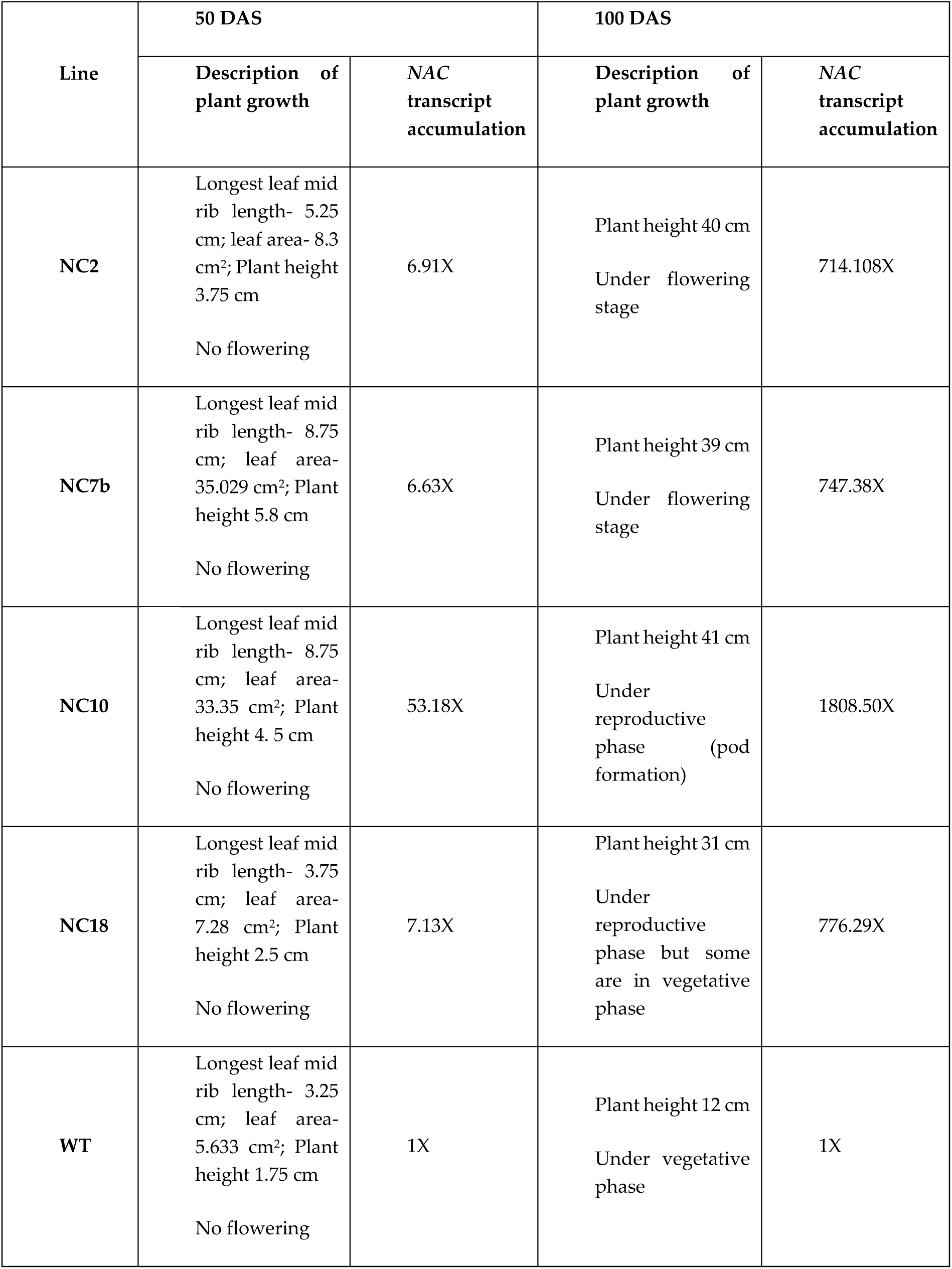
Analysis of growth and *NAC* transcript accumulation in leaf tissues in NAC lines and WT

**Table S2.** Identification of cis-acting regulatory elements for the genes being upregulated in NAC transgenic as compared to wild type

| 50. Gene Id | 51. Cis-acting regulatory elements (CAREs) |
| --- | --- |
| 52. KEH33843 | 53. ABRE, CCAAT-box, G-box, MBS, MSA-like, Sp1, TCCACCT-motif, box II, chs-Unit 1 m1 |
| 54. KEH15834 | 55. TC-rich repeats, circadian, TC-rich repeats, ACA-motif, TATA-box, CAAT-box |
| 56. XP_003616288 | 57. 3-AF1 binding site, 5UTR Py-rich stretch, AAGAA-motif, ATC-motif, ATCT-motif, AuxRR-core, Box 4, Box I, CAAT-box, ERE, GA-motif, GARE-motif, GATA-motif, GCN4_motif, GT1-motif, I-box, P-box, Sp1, TATA-box, TATCCAT/C-motif, TC-rich repeats, TCA-element, TCT-motif, as-2-box, circadian |
| 58. XP_003626838 | 59. 5UTR Py-rich stretch, AE-box, Box I, CAAT-box, ERE, GAG-motif, GATA-motif, I-box, P-box, TATA-box, TC-rich repeats, TCT-motif, circadian, rbcS-CMA7a |
| 60. XP_003613921 | 61. 5UTR Py-rich stretch, AAGAA-motif, AE-box, Box I, CAAT-box, CATT-motif, CTAG-motif, GA-motif, GARE-motif, GATA-motif, O2-site, P-box, TATA-box, as-2-box, circadian |
| 62. ABE87613 | 63. 5UTR Py-rich stretch, AE-box, ARE, CAAT-box, AT-rich element, GA-motif, GCN4-motif, P-box, TATA-box, TC-rich repeats, TCA-element, TCT-motif |
| 64. XP_009777082 | 65. 3-AF1 binding site, 5UTR Py-rich stretch, AAGAA-motif, AE-box, ARE, ATCT-motif, AuxRR-core, Box 4, CAAT-box, GARE-motif, GAG- motif, GCN4-motif, O2-site, TATA-box, TC-rich repeats, TCA-element, circadian |
| 66. CCA65997 | 67. 5UTR Py-rich stretch, AAGAA-motif, AE-box, ATCT-motif, CAAT-box, GARE-motif, GCN4-motif, Box I, P-box, TATA-box, TC-rich repeats, TCA-element, TCT-motif, as-2-box, circadian, Sp1 |
| 68. KFK39764 | 69. 5UTR Py-rich stretch, AAGAA-motif, AE-box, ATCT-motif, CAAT-box, CATT-motif, GA-motif, GAG-motif, GATA-motif, I-box, P-box, TATA-box, O2-site, TC-rich repeats, TCA-element, TCT-motif, circadian |
| 70. BAB09923 | 71. AAGAA-motif, AE-box, ARE, ATCT-motif, Box 4, CAAT-box, CATT-motif, CTAG-motif, GARE-motif, GAG- motif, GATA-motif, I-box, Skn-1_motif, TATA-box, TC-rich repeats, TCA-element, TCT-motif, circadian |
| 72. ABR01162 | 73. 3-AF1 binding site, 5UTR Py-rich stretch, AAGAA-motif, ARE, AT-rich repeats, Box 4, Box I, CAAT-box, GARE-motif, GA-motif, GT1-motif, I-box, P-box, TATA-box, O2-site, Skn-1_motif, Sp1, TC-rich repeats, TCA-element, TCT-motif, circadian, as-2-box |
| 74. WP_011875844 | 75. AAGAA-motif, CAAT-box, GAG- motif, GATA-motif, GATT-motif, MRE, Skn-1_motif, TATA-box, P-box, TCCC-motif, circadian |
| 76. WP_028229025 | 77. AAGAA-motif, Box 4, Box I, CAAT-box, CTAG-motif, F-box, GA-motif, TATA-box, TATCCAT/C-motif |
| 78. XP_009608119 | 79. 5UTR Py-rich stretch, AAGAA-motif, ATCT-motif, CAAT-box, CATT-motif, GAG- motif, GATA-motif, I-box, P-box, TATA-box, TC-rich repeats, TCA-element, circadian, as-2-box |
| 80. XP_009797331 | 81. AAGAA-motif, AE-box, CAAT-box, CTAG-motif, GAG- motif, GATA-motif, Gap-box, I- box, P-box, TATA-box, TC-rich repeats, TCA-element, WUN-motif, circadian, as-2-box |
| 82. XP_009607952 | 83. AE-box, ATCT-motif, Box 4, Box I, CAAT-box, CATT-motif, GAG-motif, GATT-motif, I-box, P-box, TATA-box, TCA-element, TC-rich repeats, circadian, as-2-box |
| 84. XP_009601984 | 85. 3-AF1 binding site, 5UTR Py-rich stretch, AAGAA-motif, AE-box, ATCT-motif, CAAT- box, GARE-motif, GA-motif, GATA-motif, GT1-motif, GCN4-motif, P-box, TATA-box, TATC-box, TCA-element, TC-rich repeats, TCT-motif, circadian |
| 86. XP_009611343 | 87. 5UTR Py-rich stretch, CAAT-box, GATA-motif, CTAG-motif, TATA-box, TCT-motif, circadian, as-2-box |
| 88. XP_009610696 | 89. AAGAA-motif, CAAT-box, GARE-motif, GAG-motif, Box 4, TATA-box, TATCCAT/C- motif, TCA-element, TC-rich repeats, circadian, as-2-box |
| 90. XP_009631607<br>91. | 92. 5UTR Py-rich stretch, AAGAA-motif, AE-box, AuxRR-core, Box 4, CAAT-box, CATT- motif, GARE-motif, HSE, O2-site, I-box, TATA-box, Skn-1_motif, TCA-element, TC-rich repeats, TCCC-motif, TCT-motif, circadian |
| 93. XP_009589281 | 94. 5UTR Py-rich stretch, AAGAA-motif, ARE, ACE, AuxRR-core, Box I, CAAT-box, CATT- motif, GARE-motif, F-box, I-box, GAG-motif, TATA-box, Skn-1_motif, TATCCAT/C-motif, TC-rich repeats, circadian, as-2-box, box E |
| 95. XP_009759366 | 96. 3-AF1 binding site, AE-box, CAAT-box, CATT-motif, I-box, Sp1, TATA-box, TCA-element, TC-rich repeats, circadian |
| 97. XP_006359484 | 98. AE-box, Box 4, CAAT-box, CATT-motif, HSE, I-box, MRE, O2-site, Skn-1_motif, TATA-box, TC-rich repeats, TCT-motif, WUN-motif, circadian |
| 99. XP_009792488 | 100. AAGAA-motif, AE-box, Box 4, CAAT-box, I-box, GAG-motif, O2-site, TATA-box, TC-rich repeats, circadian, as-2-box, box E |
| 101. XP_009593886 | 102. ARE, CAAT-box, GA-motif, GAG-motif, GARE-motif, GATA-motif, I-box, P-box, TATA- box, circadian |
| 103. XP_010520610 | 104. AAGAA-motif, AE-box, ATCT-motif, GARE-motif, GATA-motif, CAAT-box, I-box, Skn- 1_motif, GT1-motif, TATA-box, TC-rich repeats, TCA-element, TCT-motif, circadian |
| 105. XP_009627032 | 106. AE-box, ATCT-motif, CAAT-box, GA-motif, GATA-motif, GT1-motif, GCN4-motif, I-box, HSE, TATA-box, TC-rich repeats, TCA-element, TCT-motif, circadian, as-2-box |
| 107. XP_009779356 | 108. 5UTR Py-rich stretch, AAGAA-motif, Box 4, CAAT-box, GA-motif, GCN4-motif, CATT- motif, HSE, I-box, Sp1, TATA-box, TATCCAT/C-motif, TC-rich repeats, TCA-element, TCT- motif, circadian, as-2-box |
| 109. XP_009802473 | 110. 3-AF1 binding site, AAGAA-motif, ATGCAAAT motif, CAAT-box, CAT-box, GARE-motif, GCN4-motif, Skn-1_motif, O2-site, TATA-box, TCT-motif, circadian |
| 111. XP_009607141 | 112. AAGAA-motif, CAAT-box, GAG-motif, GATA-motif, Skn-1_motif, I-box, TATA-box, TCA-element |
| 113. XP_009775835 | 114. AAGAA-motif, CAAT-box, Skn-1_motif, I-box, TATA-box, TC-rich repeats, circadian |
| 115. XP_010519403 | 116. 3-AF1 binding site, 5UTR Py-rich stretch, AAGAA-motif, AE-box, CAAT-box, CATT-motif, GA-motif, GAG-motif, CTAG-motif, GATA-motif, GT1-motif, GARE-motif, I-box, Skn-1_motif, O2-site, Sp1, P-box, TATA-box, TATC-box, TC-rich repeats, TCCC-motif, TCT-motif, chs-CMA2a, circadian |
| 117. XP_010696413 | 118. AAAC-motif, AAGAA-motif, ARE, ATCT-motif, AuxRR-core, CAAT-box, CTAG-motif, GAG-motif, GARE-motif, GCN4-motif, Skn-1_motif, TATA-box, TC-rich repeats, TCA-element, circadian, as-2-box |
| 119. XP_009629427 | 120. ATGCAAAT motif, CAAT-box, Box 4, CATT-motif, G-box, GAG-motif, GARE-motif, I-box, MRE, Skn-1_motif, P-box, TATA-box, TC-rich repeats, TCA-element, TCCC-motif, TCT-motif, circadian |
| 121. XP_009785642 | 122. ABRE, CAT-box, CCAAT-box, G-Box, G-box, GATA-motif, GC-motif, MBS, MSA-like, Sp1, TCA-element, TCCACCT-motif, TGA-element, box II, circadian, plant_AP-2-like, rbcS-CMA7a |
| 123. XP_009612935 | 124. 3-AF1 binding site, 5UTR Py-rich stretch, AAGAA-motif, AE-box, ATCT-motif, ARE, ATGCAAAT motif, Box 4, CAAT-box, GA-motif, GAG-motif, CTAG-motif, GATA-motif, I-box, O2-site, TATA-box, TC-rich repeats, TCT-motif, circadian, as-2-box |
| 125. XP_009622691 | 126. 5UTR Py-rich stretch, AAGAA-motif, AE-box, ATCT-motif, CAAT-box, CATT-motif, GARE-motif, GCN4-motif, GA-motif, GT1-motif, MRE, Skn-1_motif, P-box, TATA-box, TC-rich repeats, TCA-element, TCT-motif, circadian, as-2-box, chs-CMA2a |
| 127. XP_009624162 | 128. 5UTR Py-rich stretch, AAGAA-motif, ATCT-motif, Box 4, CAAT-box, CATT-motif, GARE-motif, GATA-motif, I-box, O2-site, TATA-box, TC-rich repeats, TCA-element, TCT-motif, circadian |
| 129. XP_006360870 | 130. 3-AF1 binding site, 5UTR Py-rich stretch, AAGAA-motif, AE-box, ATCT-motif, ARE, CAAT-box, GA-motif, Box I, GAG-motif, HSE, Skn-1_motif, P-box, Sp1, TATA-box, TC-rich repeats, TCT-motif, box E, circadian |
| 131. XP_009607401 | 132. 5UTR Py-rich stretch, AE-box, AuxRR-core, Box I, CAAT-box, CATT-motif, GAG-motif, I-box, TATA-box, TCA-element, TCT-motif, circadian |
| 133. XP_009770604 | 134. 5UTR Py-rich stretch, AE-box, AAGAA-motif, Box I, Box 4, CAAT-box, CATT-motif, F-box, G-Box, GCN4-motif, I-box, P-box, Skn-1_motif, TATA-box, TC-rich repeats, TCA-element, TCT-motif, circadian |
| 135. XP_009785514 | 136. 5UTR Py-rich stretch, AAGAA-motif, AC-II, AuxRR-core, Box 4, CAAT-box, GAG-motif, GARE-motif, GATA-motif, GT1-motif, HD-Zip 3, I-box, P-box, TATA-box, TC-rich repeats, TCA-element, TCT-motif, chs-CMA2a, circadian |
| 137. WP_028229032 | 138. 5UTR Py-rich stretch, AAGAA-motif, AuxRR-core, Box 4, CAAT-box, GATA-motif, GA-motif, GT1-motif, HSE, I-box, L-box, P-box, TATA-box, TC-rich repeats, TCA-element, TCT-motif, as-2-box, circadian |
| 139. XP_009631610 | 140. 5UTR Py-rich stretch, AAGAA-motif, AE-box, Box I, CAAT-box, CTAG-motif, GA-motif, HD-Zip 3, HSE, I-box, P-box, Skn-1_motif, TATA-box, TATCCAT/C-motif, TC-rich repeats, TCA-element, circadian |
| 141. XP_009787092 | 142. 5UTR Py-rich stretch, AAGAA-motif, AE-box, Box I, CAAT-box, ATCT-motif, GAG-motif, GA-motif, GATA-motif, GCN4-motif, GT1-motif, I-box, Skn-1_motif, TATA-box, TC-rich repeats, circadian |
| 143. XP_009801279 | 144. 3-AF1 binding site, AAGAA-motif, ABRE, ARE, ATCT-motif, Box 4, CAAT-box, F-box, GATA-motif, GARE-motif, HSE, I-box, TATA-box, TATC-box, TCA-element, TCT-motif, circadian |
| 145. XP_009594285 | 146. 5UTR Py-rich stretch, AAGAA-motif, ARE, Box I, CAAT-box, CTAG-motif, ERE, GAG-motif, GARE-motif, GA-motif, Skn-1_motif, P-box, Sp1, TATA-box, TC-rich repeats, TCA-element, as-2-box, circadian |
| 147. YP_173432 | 148. 3-AF1 binding site, 5UTR Py-rich stretch, AAGAA-motif, CAAT-box, TATA-box, TCT-motif, chs-CMA2a, circadian |

## References

1. Lindemose, S.; O’Shea, C.; Jensen, M.K.; Skriver, K. Structure, function and networks of transcription factors involved in abiotic stress responses. Int. J. Mol. Sci. 2013, 14, 5842–5878.

2. Shao, H.; Wang, H.; Tang, X. NAC transcription factors in plant multiple abiotic stress responses: Progress and prospects. Front. Plant Sci. 2015, 6, 902.

3. Baillo, E.H.; Kimotho, R.N.; Zhang, Z.; Xu, P. Transcription Factors Associated with Abiotic and Biotic Stress Tolerance and Their Potential for Crops Improvement. Genes 2019, 10, 771.

4. Lata, C.; Yadav, A.; Prasad, M. Role of plant transcription factors in abiotic stress tolerance. In: Shanker A (Ed.) Abiotic stress response in plants - physiological, biochemical and genetic perspectives. InTech, New Delhi, India 2011, pp 269-296 (online only)

5. Ernst, H.A.; Olsen, A.N.; Larsen, S.; Lo Leggio. L. Structure of the conserved domain of ANAC, a member of the NAC family of transcription factors. EMBO Rep. 2004, 5, 297–303.

6. Hu, H.; You, J.; Fang, Y.; Zhu, X.; Qi, Z.; Xiong, L. Characterization of transcription factor gene SNAC2 conferring cold and salt tolerance in rice. Plant Mol. Biol. 2008, 67, 169–181.

7. Hao, Y.J.; Wei, W.; Song, Q.X.; Chen, H.W.; Zhang, Y.Q.; Wang, F.; Zou, H.F.; Lei, G.; Tian, A.G.; Zhang, W.K.; Ma, B.; Zhang, J.S.; Chen, S.Y. Soybean NAC transcription factors promote abiotic stress tolerance and lateral root formation in transgenic plants. Plant J. 2011, 68, 302–313.

8. Singh, S.; Khalid, H.; Grover, A.; Singh, A.; Nasim, M. Altered Physiological Responses of NAM, ATAF1/2 and CUC2 (NAC) Gene of *Lepidium latifolium (LlaNAC*) Over-expressing Tobacco Plants. Acta Physiol. Plant 2019, 41, 139.

9. Grover, A.; Singh, S.; Pandey, P.; Patade, V.Y.; Gupta, S.M.; Nasim, M. Overexpression of NAC gene from Lepidium latifolium enhances biomass, shortens life cycle and induces cold stress tolerance in tobacco: potential for engineering fourth generation biofuel crops. Mol. Biol. Rep. 2014, 11, 7479–7489.

10. Xu, B.; Ohtani, M.; Yamaguchi, M.; Toyooka K.; Wakazaki, M.; Sato, M.; Kubo, M.; Nakano, Y.; Sano, R.; Hiwatashi, Y.; Murata, T.; Yoneda, A.; Kato, K.; Hasebe, M.; Demura, T. Contribution of NAC transcription factors to plant adaptation to land. Science 2014, 343, 1505–1508.

11. Podzimska-Sroka, D.; O’Shea, C.; Gregersen, P.L.; Skriver, K. NAC transcription factors in senescence: From molecular structure to function in crops. Plants 2015, 4, 412–448.

12. Kim, H.J.; Nam, H.G.; Lim, P.O. Regulatory network of NAC transcription factors in leaf senescence. Curr. Opin. Plant. Biol. 2016, 33, 48–56.

13. Singh, S.; Grover, A.; Nasim, M. Biofuel potential of plants transformed genetically with NAC family genes. Front. Plant Sci. 2016, 7, 22.

14. Shamimuzzaman, M.; Vodkin, L. Genome-wide identification of binding sites for NAC and YABBY transcription factors and co-regulated genes during soybean seedling development by ChIP-Seq and RNA-Seq. BMC Genomics 2013, 14, 477.

15. Wang, W.; Yuan Y.; Yang, C.; Geng, S.; Sun, Q.; Long, L.; Cai, C.; Chu, Z.; Liu, X.; Wang, G.; Du, X.; Miao, C.; Zhang, X.; Cai, Y. Characterization, expression, and functional analysis of a novel NAC gene associated with resistance to verticillium wilt and abiotic stress in cotton. G3 (Bethesda) 2016, 6, 3951–3961.

16. Zheng, X.; Tang, S.; Zhu, S.; Dai, Q.; Liu, T. Identification of a NAC transcription factor family by deep transcriptome sequencing in onion (*Allium cepa* L.). PLoS ONE 2016, 11, e0157871.

17. Zhong, R.; Yuan, Y.; Spiekerman, J.J.; Guley, J.T.; Egbosiuba, J.C.; Ye, Z-H. Functional characterization of NAC and MYB transcription factors involved in regulation of biomass production in Switchgrass (*Panicum virgatum*). PLoS ONE 2015 10, e0134611.

18. Ordiz, M.I.; Barbas, C.F.; Beachy RN. Regulation of transgene expression in plants with polydactyl zinc finger transcription factors. Proc. Natl. Acad. Sci. U S A. 2002, 99, 13290–5.

19. Yan, L.; Loukoianov, A.; Tranquilli, G.; Helguera, M.; Fahima, T.; Dubcovsky, J. Positional cloning of the wheat vernalization gene *VRN1*. Proc. Natl. Acad. Sci. USA. 2003, 100, 6263–6268.

20. Chou, K-C.; Shen, H-B. Plant-mPLoc: a top-down strategy to augment the power for predicting plant protein subcellular localization. PLoS ONE 2010, 5, e11335.

21. Pérez-Rodríguez, P., Riaño-Pachón, D.M., Corrêa, L.G., Rensing, S.A., Kersten, B., Mueller-Roeber, B. PlnTFDB: updated content and new features of the plant transcription factor database. Nucleic Acids Res. 2010, 38, D822–D827.

22. Lescot, M.; Déhais, P.; Thijs, G.; Marchal, K.; Moreau, Y.; Van de Peer Y.; Rouzé P.; Rombauts S. PlantCARE: a database of plant cis-acting regulatory elements and a portal to tools for in silico analysis of promoter sequences. Nucleic Acids Res. 2002, 30, 325–327.

23. Aslam, M.; Grover, A.; Sinha, V.B.; Fakher, B.; Pande, V.; Patade, P.V.; Gupta, S.M.; Anandhan, S, Ahmed, Z. Isolation and characterization of cold responsive NAC gene from *Lepidium latifolium*. Mol. Biol. Rep. 2012, 39, 9629–9638.

24. Barre, A.; Bourne, Y.; Van Damme, E.J.; Peumans, W.J.; Rougé, P. Mannose-binding plant lectins: Different structural scaffolds for a common sugar-recognition process. Biochimie 2001, 83, 645–651.

25. Himmel, M.E.; Ding, S-Y.; Johnson, D.K.; Adney, W.S.; Nimlos, M.R.; Brady, J.W.; Foust, D. Biomass recalcitrance: Engineering plants and enzymes for biofuels production. Science. 2007, 315, 804–807.

26. Kleczkowski, L.A.; Kunz, S.; Wilczynska. Mechanisms of UDP-glucose synthesis in plants. Crit. Rev. Plant. Sci. 2010, 29, 191–203.

27. Kotake, T.; Hirosawa, C.; Ando, Y.; Tsumuraya Y. Generation of nucleotide sugars for biomass formation in plants. Plant Biotechnol. 2010, 27, 231–236.

28. Kleczkowski, L.A.; Decker, D.; Wilczynska, M. UDP-sugar pyrophosphorylase: a new old mechanism for sugar activation. Plant Physiol. 2011, 156, 3–10.

29. arkowski, L.P.; Van den Ende, W. Cold tolerance triggered by soluble sugars: a multifaceted countermeasure. Front. Plant Sci. 2015, 6, 203.

30. Olsen, A.N.; Ernst, H.A.; Leggio, L.L.; Skriver, K. NAC transcription factors: Structurally distinct, functionally diverse. Trends Plant Sci. 2005, 10, 79–87.

31. Hu, W.; Wei, Y.; Xia, Z.; Yan, Y.; Hou, X.; Zou, M.; Lu, C.; Wang, W.; Peng, M. Genome-wide identification and expression analysis of the NAC transcription factor family in cassava. PLoS ONE 2015, 10, e0136993.

32. Samad, A.F.A.; Sajad, M.; Nazaruddin, N.; Fauzi, I.A.; Murad, A.M.A.; Zainal, Z.; Ismail, I. MicroRNA and transcription factor: Key players in plant regulatory network. Front. Plant Sci. 2017, 8, 565.

